# Modeling Systemic Colorimetric Parameters as a Tool for Processing Images of Clumps of Toxic Cyanobacteria Targeted at Their Boundaries Detection

**DOI:** 10.1101/232413

**Authors:** Yu. G. Bespalov, K. V. Nosov, P. S. Kabalyants

## Abstract

Global climate change, along with other large-scale consequences of human impact upon the nature, increases the risk of biosafety threats associated with the disturbance of stability of communities of living organisms. In this regard, the topicality of the challenge of developing methods for monitoring and correcting homeostasis mechanisms that can support this stability is a problem of premium importance.

The work aims at investigation of techniques of remote detection of toxic cyanobacteria clumps in water area, with the use of dynamical modeling.

## Introduction

Global climate change regardless they causes — variations in solar radiation, planet tectonics, or global warning — may lead to the risk of biosafety threats associated with the lack of stability of living organisms’ communities. So, the problem of developing techniques for monitoring and correcting homeostasis mechanisms supporting this stability in communities has premium importance. As to the causes of global climate change, there is no fully unambiguous point of view among environmentalists, but in any case, there is no doubt that it is necessary to overcome their negative, sometimes dramatic, consequences. Among such consequences is a mass development of toxic cyanobacteria in fresh and marine waters [1, 2, 3].

A resonance example of the threat of biosafety for potable water consumption, in the case of implementation of such a form of stability’s disturbance in water ecosystem, is the situation in the Middle East’s Lake Kinneret [4]. Another example of such the form of instability of aquatic ecosystem is the mass development of cyanobacteria, which has been observed for several years in the Baltic Sea [5].

In this connection, a number of measures were proposed and used to correct the mechanisms of homeostasis of the aquatic ecosystem with the aim of eliminating the real or potential threat of mass development of cyanobacteria. First, we keep in mind the limitation of intake for biogenic elements (mainly, nitrogen and phosphorus) in the form of nutrients for photosynthetic organisms into reservoirs. In many cases, these measures are significantly expensive and cause restrictions for various types of business activity (as a rule, agricultural sector related).

In this respect, it is needed to conduct the measures for monitoring the state of aquatic ecosystems for optimization of the correction schemes for this state. Implementing such measures, e.g., in the case of Lake Kinneret, large-scale and costly long-term studies of various aspects of eutrophication were required [6]. We bear in mind the balance of biogenic elements, the hydrological regime and the relationships of different components of the aquatic ecosystem.

The process of eutrophication creating the real and potential threats of mass development of cyanobacteria develops on vast water and catchment areas. Nowadays, remote (aerospace) methods are widely used to control these processes [7]. In [8], a number of difficulties is mentioned that arises when solving the problem of determining cyanobacterial accumulations in marine waters by analyzing satellite images.

These problems will have a different nature for extreme situations of emergence of mutants of cyanobacteria, the toxins of which will create biosafety threats that may be more serious than those created by natural strains of cyanobacteria currently known. (Unfortunately, there is a real danger of appearance of such mutants: in natural way or as a result of activities of bioterrorist organizations).

In such extreme situations, the use of algicides for destruction of cyanobacterial clumps, in which such new especially dangerous toxins will be produced, may have sense.

The solution of this problem is also necessary for planning and implementation of biosafety measures in the coastal areas which cyanobacterial clumps move to.

In such extreme situations, in conditions of lack of time, it may be necessary to mobilize all available tools for remote sensing water areas. They may include, in particular, light drones with relatively simple and inexpensive equipment for digital photography on their board. This equipment enables to take photos only in the visible band of the spectrum and record the brightness of the photographed objects in wide spectral bands corresponding to the red, green, and blue components.

Earlier [9], the possibility of description for some aspects of the dynamics of colorimetric parameters of plant communities and protective coloration of animals was shown on the basis of such initial factual information obtained by mrelatively crude techniques. This possibility is available with the use of discrete models of dynamical systems (DMDS) already applied to describing the structure of relationships and dynamics of components of systems of quite different nature [10]

A DMDS model provide the explicit description of the structure of between-component relationships, which express pairwise positive and negative between-component influences, while intra-component influences are symmetric. The relations of this type is an “alphabet” of between-species or between-organisms interaction in theoretical biology [11,12, 13]. Using the structure of relationships between all the system’s components and given initial conditions of the system, a trajectory of the system can be calculated. Taking into account a finite number of the system’s states, the trajectory will be periodical. Such the trajectory is called an idealized system trajectory (ITS) and, hence, it presents a periodical cycle of changes in values of all the components of the system. These values are measured semi-quantitatively, in conditional scores.

In a number of cases, such a description makes it possible to solve the problem of identifying a given type of plant communities according to the nature of systemic aspects of the dynamics of above-mentioned relatively crude colorimetric parameters associated with the content of all forms of chlorophyll and all types of carotenoids and other yellow-orange plant pigments. Considering these systemic aspects, it is possible to introduce systemic colorimetric parameters, SCPs, the use of which in the processing of images of water areas will enable to determine the localization and boundaries of cyanobacterial clumps (bloom patches).

The values of the SCPs should be determined on the basis of the data on the values of primary colorimetric parameters (PCPs), which can be obtained by computer analysis of components of the RGB image model. Such a technique provides comparatively crude initial factual data, but from the viewpoint of reducing financial costs and the possibility of mobilizing a wider arsenal of remote data collection facilities in extreme situations, has a number of advantages over more exact remote methods based on registration of brightness in narrow spectral bands, in particular, used for detecting clumps of cyanobacteria [13].

The present work aims at investigation of possibilities of detection of clumps of toxic cyanobacteria (bloom patches) in the water area, with the use of DMDS modeling. We keep in mind detection of localization and boundaries of bloom patches by processing the images obtained by digital photography in the visible spectral range, with the selection of spectral bands corresponding to the components of the RGB model of digital photography, using SCPs.

In this work, a draft version of such a technique is proposed.

## Material and methods

For drawing working hypotheses regarding the type of SCPs, we have analyzed the ITS earlier calculated and described in [15]. This ITS describes the dynamics of the PCPs for a bloom patch, which is a primitive plant community. This ITS is presented in Fig. 1.

**Fig. 1.**
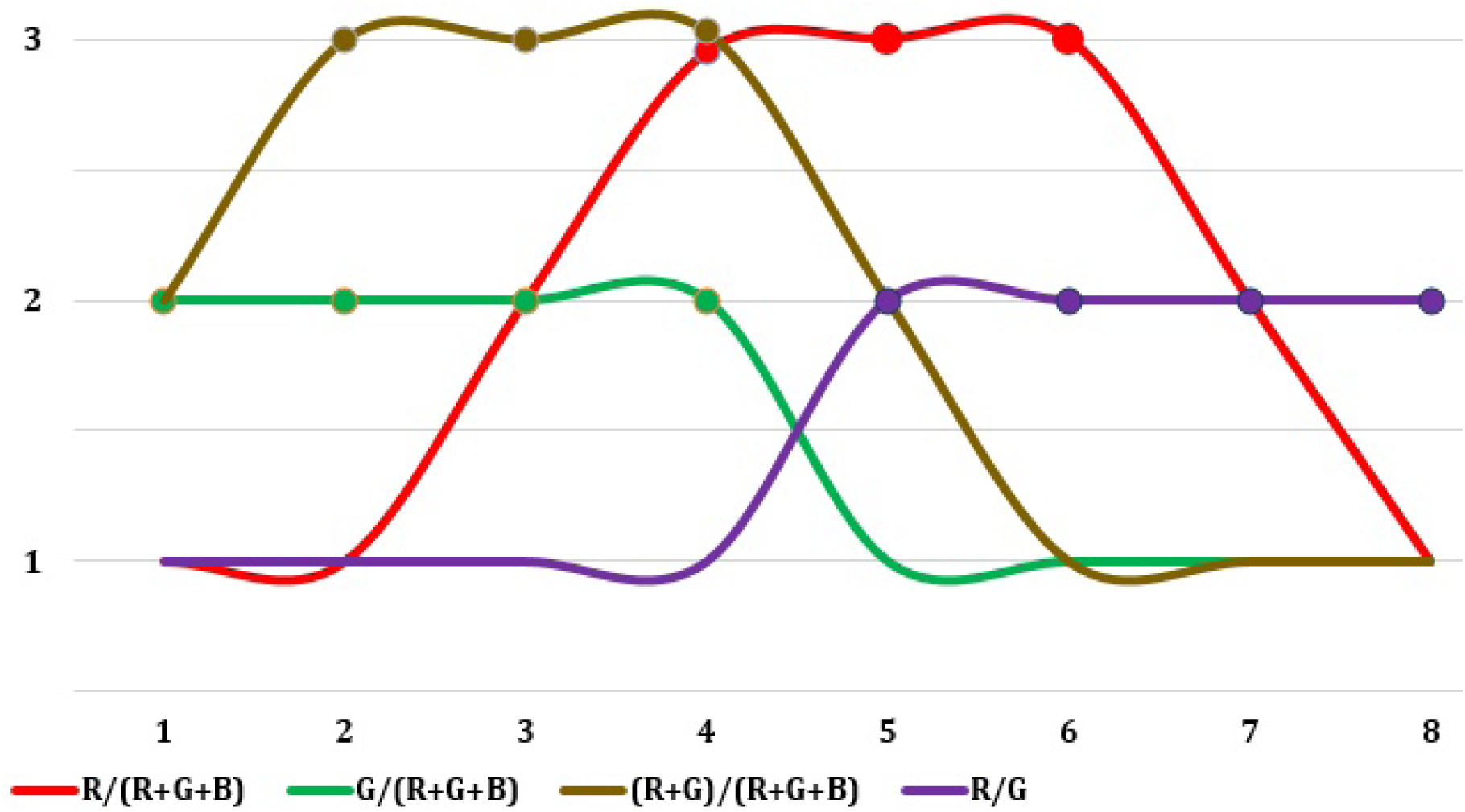
Idealized trajectory of the system describing the dynamics of the values of the primary colorimetric parameters of the cyanobacteria community. Smooth curves are used for better visual perception. The levels of values are measured in conditional scores and mean: 1 - low, 2 - medium, 3 - high. X axis: time, Y axis: the values of colorimetric parameters. The maximum values of each colorimetric parameter in the ITS are marked by round markers.

The mean of PCPs:

R/(R+G+B) — reflects the amount of orange and yellow pigments dominated in old dying and dead plant cells;

G/(R+G+B) — reflects the amount of green pigment of chlorophyll dominated in young actively photosynthetic cells;

(R+G)/(R+G+B) — reflects the total number of all cells, young and old, dying and dead;

R/G — reflects the value of the “yellow-green index”, the indicator of pigment diversity.

For computation of this ITS, an earlier mentioned approach under the title “rechronization” that describes the dynamics of the colorimetric parameters of the cereal communities [16] was used. This approach enables to build ITSs on the basis of one snapshot of the objects, which dynamics we are modeling. The rechronization is based on the following: different parts of a plant community located on different areas of a habitat have a common cycle of changes (e.g., of their colorimetric parameters), but at any moment may be at different phases of changes. These phases correspond to different time moments in the ITS.

The working hypotheses obtained on the basis of the ITS’s analysis were verified by processing the image of Baltic Sea surface with a bloom patch.

## Statement of the problem and discussion of the results

In the course of tackling the problem, we consider a bloom patch as a primitive algae community, which has a certain cycle of states’ changes. Due to some physical and chemical properties of cyanobacteria clumps, they possess a greater ability to retain their shape and size than the suspension of algal cells in a water area outside the bloom patches. Accordingly, this cycle of states’ changes of a bloom patch has also a certain stability. Suspension of algae outside the bloom patch does not have such a cycle, its state is changing primarily due to influence of hydrological factors. (Of course, this distinction between the bloom patches and suspension of algae beyond the patches' boundaries have place only under relatively weak wind and wave that mix the water.) However, a sufficiently strong mechanical mixing of water simply destroys bloom patches.

The purpose of image processing is to detect the localization and boundaries of blooming areas. This can be achieved by the designation of different contrast colors to water areas with different variability of a certain SCP. For practical dealing this problem, a free-access Matlab package available online (https://sites.google.com/site/improctoolkit)can be used.

To perform this, one has to find a SCP with a relatively small changes of values within the cycle of changes of a bloom patch. This aspect of formulation of the problem is based on the fact that random, hydrological-related changes in relationships of a SCP within the bloom patch will be much weaker than beyond its boundaries. If we are talking about changes within the area of a bloom patch, then the role of the mentioned SCP can play, particularly, the product of the values of the two PCPs having a maximum distance (in time) between maximum values of these PCPs in the ITS. Analysis of the type of the ITS presented in Table 1 allows us to find several such pairs of control panels.

Within the framework of this paper, the product of the values of the components R/(R+G+B) and G/(R+G+B) is used as a SPC. Hereinafter, for convenience of use, we use it in simplified form R *G/(R+G+B).

For processing the image of the water area with the bloom patch, the following procedure was used. The image was divided into segments, each segment was in turn split into microsegments. For each microsegment, the average value of the expression R*G/(R+G+B) was calculated. For each segment, the value of standard deviation was calculated given the values of R*G/(R+G+B) of microsegments compose the sample. In the processed image, the value of standard deviation of each segment presents intensity of color.

The results of such the processing of the photo of Baltic Sea water area with a bloom patch is shown in Fig. 2, Fig. 3 and Fig. 4. The image in Fig. 3 was obtained from the initial image in Fig. 2 by adding computationally generated noise. This noise emulates a negative effect of poor visibility conditions, imposed, for example, by fog, atmosphere precipitations etc.

**Fig. 2.**
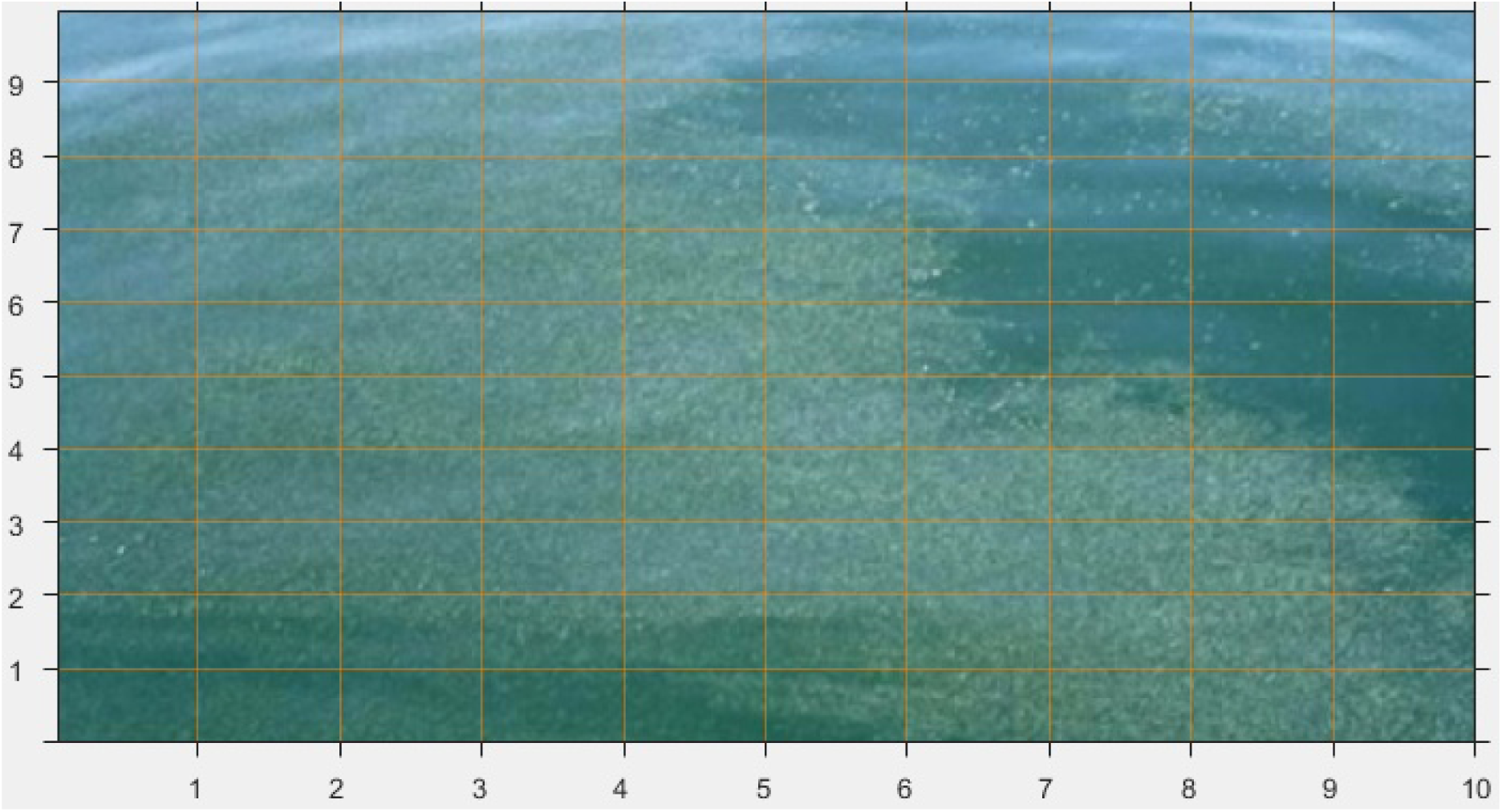
The initial image of Baltic Sea area with the bloom patch.

**Fig. 3.**
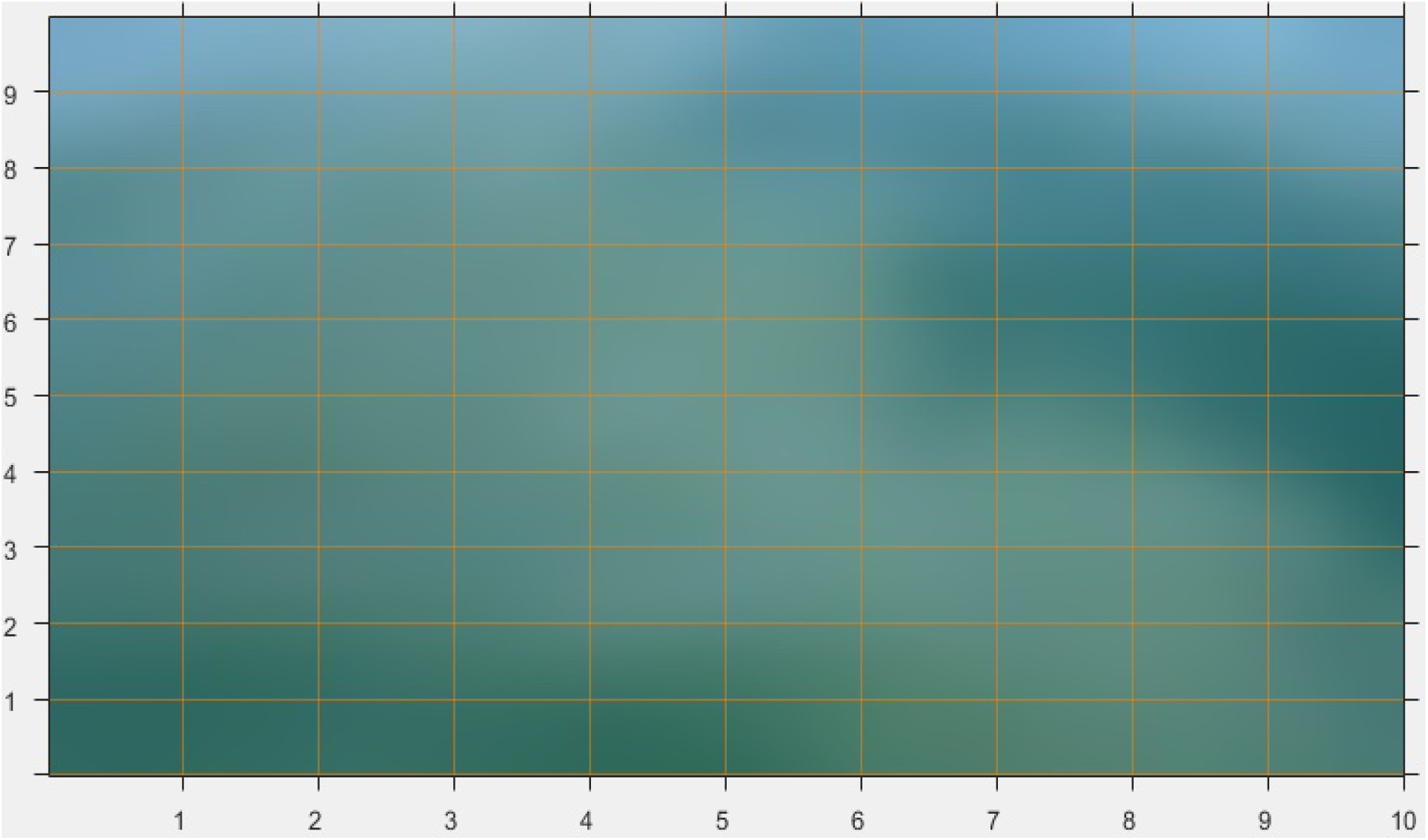
The initial image of Baltic Sea area with the bloom patch shown in Fig. 2, after the computer imitation of the poor visibility.

**Fig. 4.**
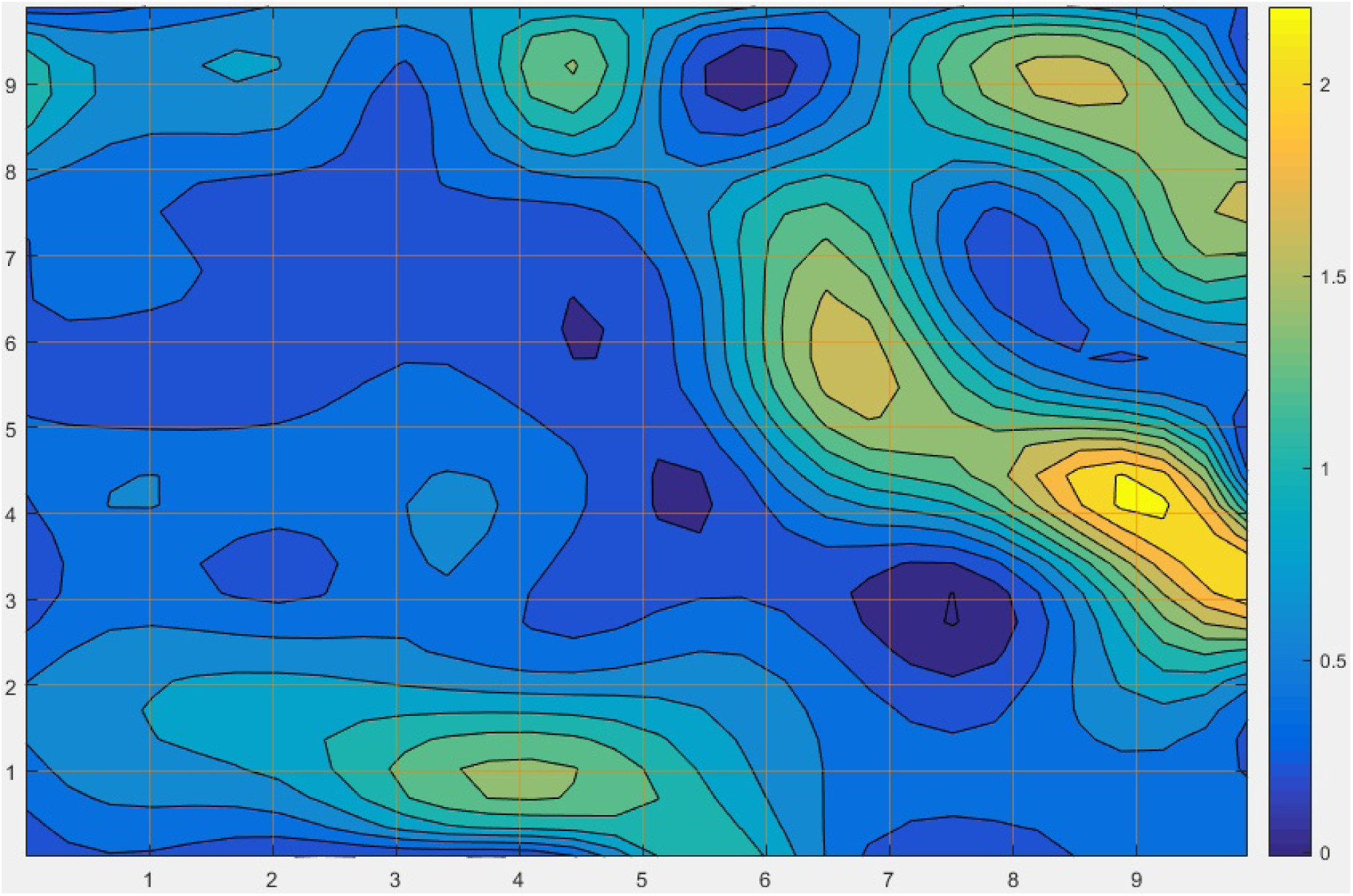
The image of the Baltic Sea area with the bloom patch after processing the noised image (Fig. 3). The image reflects the value of standard deviation of the sample related to each segment of the split. On the right is a scale of intensity corresponding to different levels of the used SPC.

The results presented above confirm the effectiveness and prospects of the approach proposed in this article for addressing the problem of processing images of the water area for detection of localization and boundaries of bloom patches.

The suggested approach differs from the known techniques used for remote identification of clumps of toxic cyanobacteria by measurements of brightness in narrow spectral band that correspond to the pigments of these photosynthetic organisms. As was mentioned, there are numerous complicated problems that arise when implementing spectral methods [8]. Besides, we should notice that such techniques require expensive complex equipment and significant financial costs for their application.

In the paper, we suggest an approach to solving the problem for identifying clumps of toxic cyanobacteria with the use of DMDS and rechronization. These techniques involve primary colorimetric parameters that can be measured by relatively simple and inexpensive equipment. The nature of this dynamics is largely determined by the prevalence of mineralization of a dead organic matter in biological communities that perform in bloom patches. In particular, this circumstance determines a low degree of coincidence of the moments with maximum values of a PCP related to green and red parts of spectrum observed in the ITS, and, accordingly, to the processes of production and dieaway of plant biomass

This regularity allowed us to suggest a specific SCP for image processing. This SCP can be considered as a reduced version of the Shannon index, which enables to estimate only equality of values and only for one pair of components of the system. The results of processing the snapshot of the water area with the bloom patch shown in Fig. 2-4, in our opinion, confirm the prospects of the approach suggested in this article.

Besides our consideration, there is a wide range of questions: the choice of the size of segments and microsegments, their allocation (in a rectangular mesh without intersection or in other way), applicability of other parameters of variability and so on. Nevertheless, the techniques proposed in the paper, in the authors' opinion, can composed a basis for methods of remote sensing regarding identification of clumps of microorganisms and algae at the surface of water area.

